# Optogenetically Evoked Accumbal Dopamine Transients Are Sufficient to Drive Locomotor Sensitization and Cross-Sensitization to Cocaine

**DOI:** 10.64898/2026.01.16.699633

**Authors:** Eva Xia Wang, Agnès Hiver, Laurena Python, Christian Lüscher, Vincent Pascoli

## Abstract

Repeated exposure to psychostimulants produces locomotor sensitization, a durable behavioral adaptation thought to reflect enhanced incentive salience driven by mesolimbic dopamine. However, the causal contribution of dopamine transients themselves, independent of drug pharmacology, remains elusive. Here we show that repeated optogenetic activation of ventral tegmental area (VTA) dopamine neurons is sufficient to induce persistent locomotor sensitization. Across successive stimulation sessions, mice exhibited a progressive escalation of locomotor activity that persisted for at least ten days after the last stimulation. Sensitization generalized beyond laser-on epochs, elevating baseline locomotion throughout the session. Importantly, mice previously exposed to optogenetic dopamine neuron stimulation displayed an enhanced locomotor response to a subsequent cocaine challenge, demonstrating cross-sensitization between optogenetic and pharmacological reinforcers. These findings establish phasic dopamine neuron activation as a sufficient driver of locomotor sensitization and reveal shared neural substrates underlying dopamine-dependent behavioral plasticity induced by optogenetic and drug reinforcers.

## Introduction

Repeated exposure to psychostimulants such as cocaine induces a progressive and persistent increase in locomotor activity, referred to as locomotor sensitization^1^. This behavioral adaptation can persist for weeks after drug withdrawal and is commonly interpreted as a manifestation of enhanced incentive salience.

Cocaine-induced locomotor sensitization requires dopamine release in the nucleus accumbens and downstream activation of D1 receptor–dependent signaling cascades, including Gα_olf_-mediated adenylate cyclase stimulation and phosphorylation of DARPP-32, engagement of NMDA receptor signaling, and ERK phosphorylation^2–7^.

At the circuit level, it is accompanied by long-lasting potentiation of medium-sized spiny neurons (MSNs) excitatory synapses arising from cortical and hippocampal inputs^8,9^. With repeated exposure to optogenetic stimulation of VTA dopamine neurons, long-lasting potentiation of these synapses underlying drug seeking was observed^10^. One injection of cocaine or brief optogenetic stimulation of VTA dopamine neurons (oDAS) drives early forms of synaptic plasticity at excitatory synapses onto DA neurons^11,12^. Thus, oDAS may recapitulate core dopamine-dependent cellular substrates of cocaine neuroadaptation while bypassing cocaine’s non-dopamine actions, but the behavioral impact on the locomotor response and sensitization has not been tested. Here we show that optogenetic VTA dopamine neuron stimulation (oDAS) is sufficient to induce locomotor sensitization and that sensitization gates an enhanced locomotor response to cocaine, arguing for cross-sensitization.

## Results

### Repeated optogenetic activation of VTA dopamine neurons induces persistent locomotor sensitization

To test whether repeated phasic activation of midbrain dopamine neurons is sufficient to induce locomotor sensitization, we repeatedly stimulated ventral tegmental area (VTA) dopamine neurons in DAT-Cre mice expressing channelrhodopsin (oDAS; Fig. 1A–C). Mice underwent daily sessions consisting of a free exploration period of the circular corridor, followed by an optogenetic stimulation period for five consecutive days, with a stimulation challenge performed ten days later in the same arena (day 15; Fig. 1B). oDAS consisted of 60 on and 60 off periods over 30 minutes (Fig. 1C, see methods).

**Figure 1.**
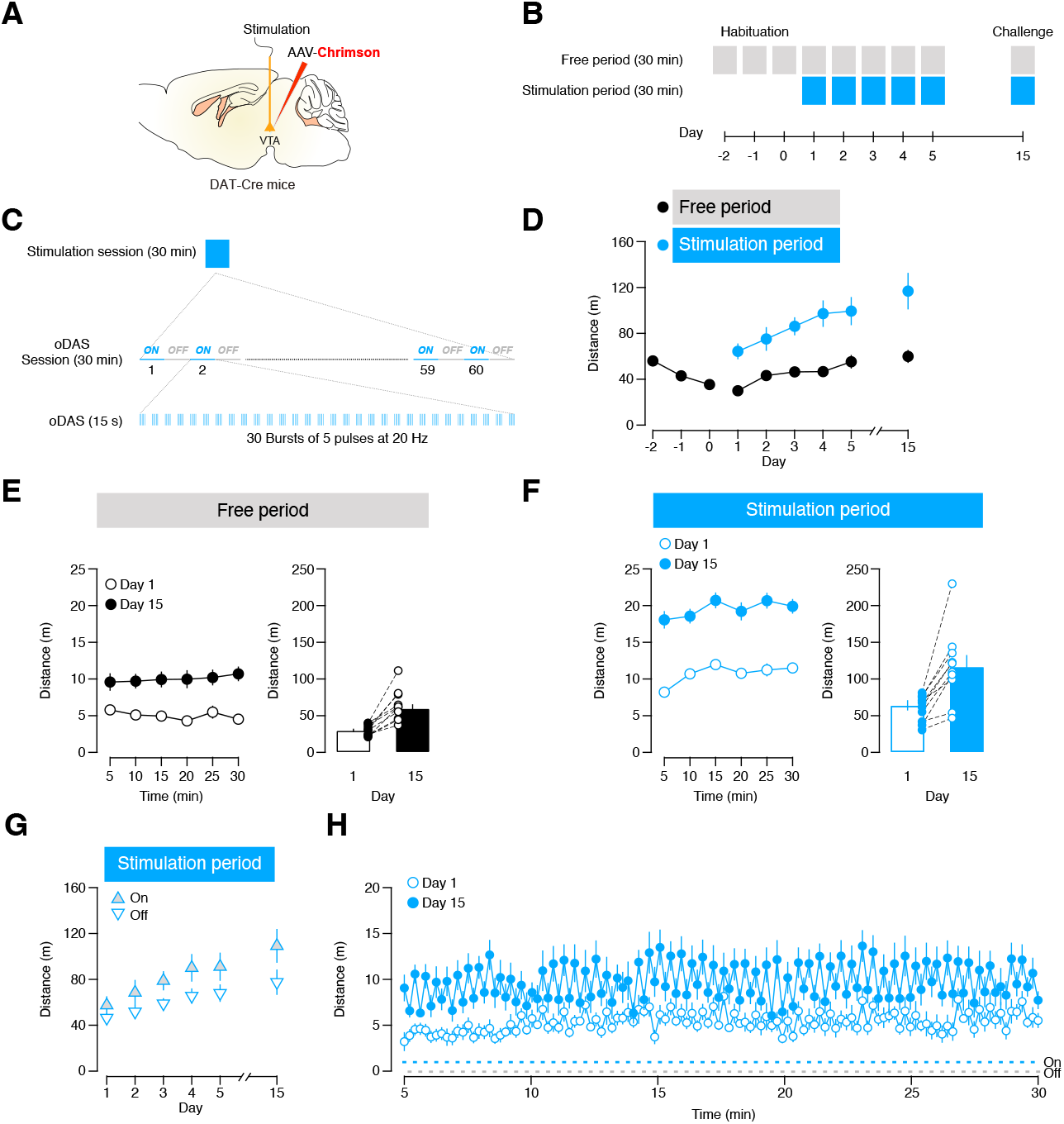
Optogenetic stimulation of VTA dopamine neurons induces locomotor sensitization to oDAS. A. mice preparation for the optogenetic stimulation of VTA dopamine neuron (oDAS). B. Experimental schedule for locomotor sessions in the circular corridor. C. oDAS protocol. D. Total distance traveled in 30 min of free periods and stimulation periods (n = 10 mice, two-way repeated measures ANOVA, period effect F_(1,18)_ = 35.04, *P* < 0.001, day effect F_(5,18)_ = 19.25, *P* < 0.001 and period x day interaction effect F_(5,18)_ = 8.32, *P* = 0.002, followed by a Bonferroni test: t_9_ = 5.00, t_9_ = 5.02, t_9_ = 6.19, t_9_ = 5.66, t_9_ = 5.34, t_9_ = 5,46, ^###^*P* < 0.001 for free vs stimulation period at day 1, day 2, day 3, day 4, day 5 and day 15, respectively; t_9_ = 4.74, t_9_ = 6.87, t_9_ = 5,49, t_9_ = 4.92, t_9_ = 6.39, ^*^*P* = 0.016, ^**^*P* = 0.001, ^**^*P* = 0.006, ^*^*P* = 0.012, ^**^*P* = 0.002 for day 1 vs day 2, day 1 vs day 3, day 1 vs day 4, day 1 vs day 5 and day 1 vs day 15, respectively, at free periods; t_9_ = 4.42, t_9_ = 6.52, t_9_ = 6.13, t_9_ = 4.74, t_9_ = 5.81, ^*^*P* = 0.025, ^**^*P* = 0.002, ^**^*P* = 0.003, ^*^*P* = 0.016, ^**^*P* = 0.004 for day 1 vs day 2, day 1 vs day 3, day 1 vs day 4, day 1 vs day 5 and day 1 vs day 15, respectively, at stimulation periods). E. Distance traveled in 30 min of free period on day 1 and day 15, in presentation of line plots per 5-min bins (two-way repeated measures ANOVA, day effect F_(1,18)_ = 40.32, *P* < 0.001, time effect F_(5,18)_ = 0.40, *P* = 0.770 and day x time interaction effect F_(5,18)_ = 0.68, *P* = 0.573, followed by a Bonferroni test: t_9_ = 3.73, t_9_ = 5.09, t_9_ = 5.71, t_9_ = 4.32, t_9_ = 3.44, t_9_ = 5.28, ^##^*P* = 0.05, ^###^*P* < 0.001, ^###^*P* < 0.001, ^##^*P* = 0.002, ^##^*P* = 0.007, ^###^*P* < 0.001 for day 1 vs day 15 at time 5, 10, 15, 25 and 30, respectively) and scatter plots of individual scores (right, paired t test, t_9_ = 6.39, ^***^*P* < 0.001). F. Distance traveled in 30 min of stimulation periods on day 1 and day 15, in presentation of line plots per 5-min bins (left, two-way repeated measures ANOVA, day effect F_(1,18)_ = 33.57, *P* < 0.001, time effect F_(5,18)_ = 8.08, *P* < 0.001 and day x time interaction effect F_(5,18)_= 0.79, *P* = 0.494, followed by a Bonferroni test: t_9_ = 8.04, t_9_ = 5.06, t_9_ = 4.24, t_9_ = 4.94, t_9_ = 5.29, t_9_ = 5.98, ^###^*P* < 0.001, ^###^*P* < 0.001, ^##^*P* = 0.002, ^###^P < 0.001, ^###^P < 0.001, ^###^P < 0.001 for day 1 vs day 15 at time 5, 10, 15, 25 and 30, respectively; t_9_ = 4.14, t_9_ = 4,17, t_9_ = 4.28, *P* = 0.038, *P* = 0.036, *P* = 0.031 for time 5 vs 10, 5 vs 25 and 5 vs 30 at day 1; t_9_ = 4.95, *P =* 0.012 for time 5 vs 30 at day 15) and scatter plots of individual scores (right, paired t test, t_9_ = 5.81, ^***^*P* < 0.001). G. Total distance traveled during oDAS-on epochs (15 min) and oDAS-off epochs (15 min) of the stimulation periods on day 1–5 and day 15 (two-way repeated measures ANOVA, laser effect F_(1,18)_ = 19.42, *P* = 0.002, main day effect F_(5,18)_ = 18.11, *P* < 0.001 and laser x day interaction effect F_(5,18)_ = 12.33, *P* = 0.001, followed by a Bonferroni test: t_9_ = 2.30, t_9_ = 2.69, t_9_ = 3.52, t_9_ = 4.06, t_9_ = 4.85, t_9_ = 9.65, ^##^*P* = 0.047, ^##^*P* = 0.025, ^##^*P* = 0.007, ^##^*P* = 0.003, ^###^*P* < 0.001, ^###^*P* < 0.001 for oDAS-on *versus* oDAS-off at day 1, day 2, day 3, day 4, day 5 and day 15, respectively; t_9_ = 4.28, t_9_ = 6.36, t_9_ = 6.40, t_9_ = 5.37, t_9_ = 7.26, ^*\**^*P* = 0.031, ^**^*P* = 0.002, ^**^*P* = 0.002, ^*\*\**^*P* = 0.007, ^***^*P* < 0.001 for day 1 *versus* day 2, day 1 *versus* day 3, day 1 *versus* day 4, day 1 *versus* day 5 and day 1 vs day 15, respectively, at oDAS-on; t_9_ = 4.00, t_9_ = 5.51, t_9_ = 5.24, t_9_ = 4.22, ^*^*P* = 0.047, ^**^*P* = 0.006, ^**^*P* = 0.008, ^*^*P* = 0.034 for day 1 *versus* day 2, day 1 *versus* day 3, day1 *versus* day 4, day1 *versus* day 15, respectively, at oDAS-off). H. Distance traveled in 60 alternating oDAS-on and oDAS-off at day 1 and day 15.

When mice first habituated to the arena, locomotion decreased over three days. Next, optogenetic stimulation produced a robust increase in locomotor activity relative to the preceding free periods starting with the first day (Fig. 1D). oDAS thus triggers an acute locomotor response. Importantly, locomotion progressively escalated across days, revealing a clear sensitization effect. Two-way repeated measures ANOVA confirmed significant main effects of period (free vs stimulation), day, and a period × day interaction, indicating that the locomotor response to oDAS increased with repeated exposure. Post hoc comparisons showed that locomotion during both free and stimulation periods was significantly elevated from day 2 onward compared with day 1, and remained significantly higher at challenge day 15 (Fig. 1D).

To assess the temporal dynamics of this effect during the 30 min sessions, we compared locomotion during the free periods on day 1. Distance traveled was significantly greater on day 15 across the entire 30-min session, with no interaction with time bin, indicating a stable upward shift rather than altered within-session dynamics (Fig. 1E). Thus, repeated oDAS induced an increase in locomotor activity starting from Day 2, in absence of the stimulation that persisted for at least ten days after the last stimulation. This increase in the locomotion during the free periods is likely due to a conditioning of the context (see below).

Locomotor activity during oDAS was significantly higher on day 15 compared with day 1 across time bins (Fig. 1F), demonstrating a persistent sensitization of the locomotor response to dopamine neuron activation itself. Remarkably, the enhanced locomotion persisted during the entire session. Together, these data show that repeated optogenetic dopamine neuron stimulation is sufficient to induce a durable form of locomotor sensitization . To further evaluate the effect of oDAS on locomotor activity we next analyzed locomotion separately during laser-on (oDAS) and laser-off epochs across days of the 30 min stimulation periods (Fig. 1G). Total distance traveled increased across days in both epochs. This indicates that sensitization generalized beyond the immediate laser-on periods and elevated locomotion throughout the session.

Consistent with this interpretation, analysis of individual stimulation cycles with high temporal resolution revealed increased locomotor activity during both laser-on and laser-off epochs on day 15 compared with day 1 (Fig. 1H). These findings indicate that repeated oDAS induces a global enhancement of locomotor drive rather than a narrowly time-locked response to optical stimulation.

### Cross-sensitization between oDAS and cocaine

To determine whether oDAS-induced sensitization share common mechanisms with psychostimulant responses, we next assessed locomotor responses to a cocaine challenge following oDAS (Fig. 2A). DAT-Cre mice and Cre-negative controls underwent 5 daily oDAS sessions as described above but then received a cocaine challenge on day 15.

**Figure 2.**
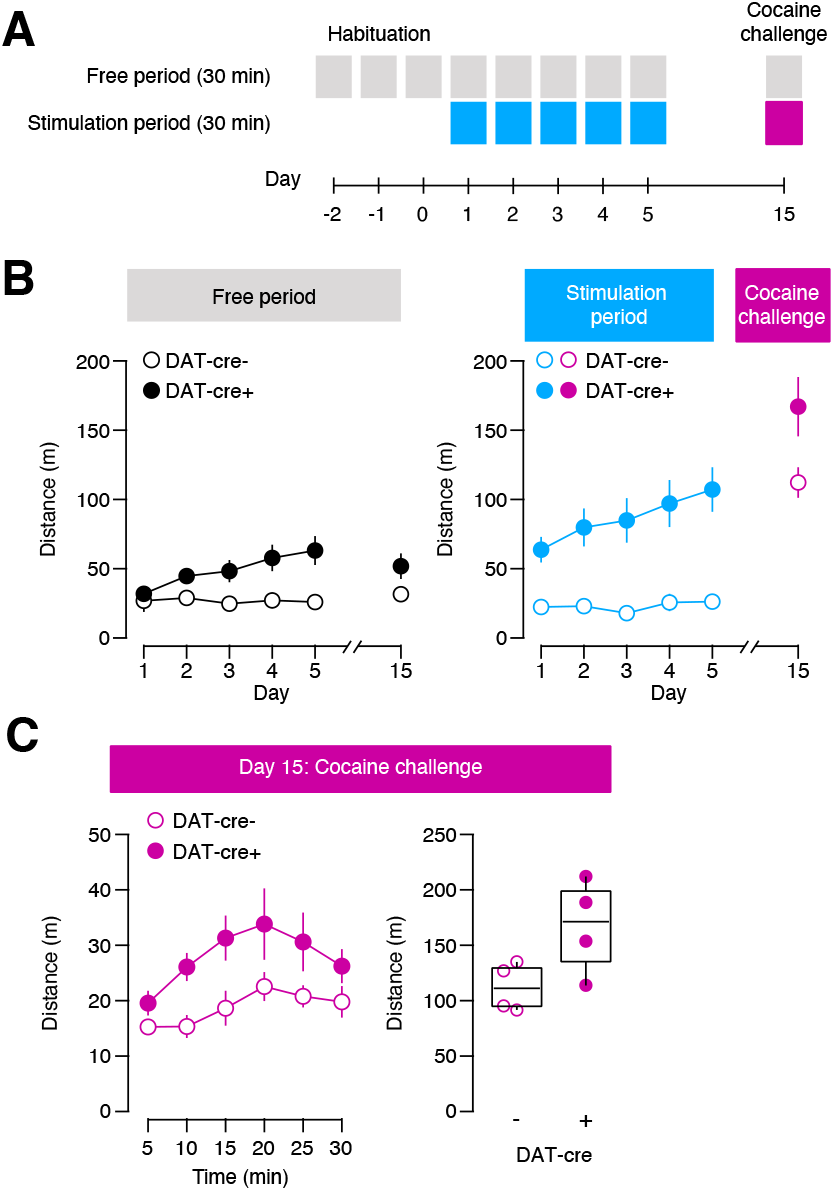
Cross-sensitization between oDAS and cocaine. A. Experimental schedule for locomotor sessions . B. Total distance traveled by DAT-Cre (n = 4 mice) and control animals (n = 4 mice) in 30 min of free period (left, two-way repeated measures ANOVA, group effect F_(1,6)_ = 9.09, *P* = 0.024, day effect F_(5,6)_ = 2.95, P = 0.086 and group x day interaction effect F_(5,6)_ = 3.06, *P* = 0.080, followed by a Bonferroni test: t_3_ = 2.73, t_3_ = 3.47, *P* = 0.042, *P* = 0.034, for DAT-Cre+ *versus* DAT-Cre-at day 4 and day 5, respectively) and stimulation and cocaine period (right, two-way repeated measures ANOVA, group effect F_(1,6)_ = 20.30, *P* = 0.004, day effect F_(5,6)_ = 41.48, *P* =< 0.001 and group x day interaction effect F_(5,6)_ = 1.61, *P* = 0.243, followed by a Bonferroni test: t_3_ = 3.93, t_3_ = 4.09, t_3_ = 4.14, t_3_ = 3.92, t_3_ = 4.91, ^#^*P* = 0.012, ^#^*P* = 0.023, ^#^*P* = 0.024, ^#^*P* = 0.019, ^#^*P* = 0.014, for DAT-Cre+ vs DAT-Cre-at day 1, day 2, day 3, day 4 and day 5, respectively). C. Distance traveled in 30 min of cocaine period by DAT-Cre and control animals in presentation of line plots per 5-min bins (left, two-way repeated measures ANOVA, group effect F_(1,6)_ = 5.21, *P* = 0.063, time effect F_(5,6)_ = 6.76, *P* = 0.005 and day x time interaction effect F_(5,6)_ = 1.13, *P* = 0.360) and scatter plots of individual scores (right, unpaired t test, t_6_ = 2.28, *P* = 0.063).

During free periods, locomotor activity was higher in DAT-Cre than in control mice (Fig. 2B, left), indicating a context conditioning as observed above . Similarly, during the stimulation period, DAT-Cre mice displayed markedly enhanced locomotor responses relative to controls mice (Fig. 2B, right panel). Finally, 10 days later, DAT-Cre+ and DAT-Cre– mice, received a challenge injection of cocaine (15 mg/kg, ip). Cocaine induced a locomotor response that was higher in DAT-Cre+ oDAS sensitized mice compared to not sensitized mice. This indicates a cross-sensitization between oDAS and cocaine.

When locomotion during the cocaine period was analyzed in 5-min bins, the response peaked 10 min after injection. DAT-cre+ mice exhibited elevated activity throughout the session compared with controls (Fig. 2C, left). This effect was robustly induced in every animal, with significantly greater total distance traveled in DAT-Cre mice (Fig. 2C, right panel). These results demonstrate cross-sensitization between repeated optogenetic dopamine neuron stimulation and cocaine-induced locomotion.

## Discussion

Optogentic stimulation induce an acute locomotor response. Across successive oDAS sessions, mice display a progressive increase in locomotor activity in response to dopamine neuron stimulation, closely resembling the locomotor sensitization observed following repeated psychostimulants administration. This enhanced locomotor response persists for at least ten days after the last stimulation session, remaining approximately doubled relative to the acute response on day 1. These effects occur in the absence of drug exposure, demonstrating that repeated phasic dopamine transients alone are sufficient to drive sensitization. Because oDAS avoids pharmacokinetic and off-target confounds, these findings establish a direct causal role for mesolimbic dopamine signaling in the induction and maintenance of locomotor sensitization.

Because oDAS and cocaine converge on shared dopamine-dependent synaptic mechanisms, this paradigm also provides a framework to investigate cross-sensitization. Our data demonstrate that oDAS sensitizes mesolimbic circuits and yields an enhanced cocaine locomotor response. Conversely, cocaine exposure may occlude oDAS-induced locomotor sensitization. In fact, in the operant condition of optogenetic DA neuron self-stimulation, i.p. cocaine injections decrease lever pressing in a dose-dependent fashion^13^. Demonstrating such bidirectional occlusion and cross-sensitization will further support the existence of overlapping neural substrates underlying dopamine-driven sensitization across optogenetic and pharmacological reinforcers. By decoupling dopamine signaling from drug-specific pharmacology, oDAS enables direct testing of which components of locomotor sensitization are attributable to dopamine transients per se, also under conditions of self-stimulation^13^.

Likewise, it will be of interest to probe whether optogenetic inhibition of VTA GABA neurons, another reinforcing optogenetic protocol^14^, can also lead to locomotor sensitization. Moreover, given the ease to control the duration of the optogenetic manipulation, it will be possible to test whether dopamine evoked synaptic plasticity in the NAc is gradual or a switch-like all-or-none phenomenon.

## Conclusions

Repeated optogenetic activation of VTA dopamine neurons via oDAS is sufficient to induce a persistent locomotor sensitization that closely parallels cocaine-induced sensitization at behavioral and synaptic levels. Future studies will have to test whether a similar cross-senitization may exist with opioid exposure^15^. These findings establish oDAS as a dopamine-specific model for studying the induction, expression, and cross-sensitization of stimulant-like behavioral adaptations, providing a controlled experimental platform to dissect the shared and distinct mechanisms underlying optogenetic and drug-induced sensitization.

## Methods

### Mice

DAT-IRES-Cre (B6.SJL-Slc6a3^tm1.1(cre)Bkmn/J^) mice and Cre-negative littermates of both sexes, aged 8–12 weeks, were from the Jackson Laboratory. On arrival, the mice were housed in groups of 3–4. All animals were kept in a temperature-(21 ± 2 °C) and hygrometry-(50 ± 5%) controlled environment with a 12 h light/12 h dark cycle, and provided with food and water *ad libitum*. Weights, sexes, and ages were distributed homogeneously among the groups . All procedures were approved by the Institutional Animal Care and Use Committee of the University of Geneva and by the animal welfare committee of the Canton of Geneva, in accordance with Swiss law.

### Virus injection and implantation

Mice were anaesthetized with a mixture of isoflurane (induction 3%, maintenance 1.5%, Attane) and O_2_ (compact anaesthesia station from Minerve) during surgery. The body temperature was maintained at 37 °C with a temperature controller system. The eyes were protected from dehydration with Lacryvisc (Alcon, Switzerland). Asepsis (betadine) and local anesthetic (Lidocaine, 0.5%) were applied before animals were placed in a stereotaxic frame (Stoeling).

AAV5-EF1a-DIO-hChR2(H134R)-mCherry (UNC) was injected in VTA (anterior–posterior, −3.28 mm; medio–lateral, −0.9 mm; dorso–ventral, −4.3 mm; at a 10° angle), in a volume of 0.5 µL, with graduated pipettes (Drummond Scientific Company) broken back to a tip diameter of 10–15 µm, at an infusion rate of 0.05 µL min^-1^. After the injection, the pipette was left in the place for 5 min to allow diffusion of the virus.

During the same surgical procedure, three screws were fixed into the skull to secure the optical implant. An optic fiber (0.2 mm diameter, Inper) was implanted 200 µm above the virus injection site and secured with dental cement. After the surgery, animals rested for 4 weeks for recovery and virus incubation.

### Optogenetic stimulation of ventral tegmental area (VTA) dopamine neuron (oDAS)

The fiber for optogenetic stimulation was connected via patch cords (BFO-1×2-F-W1.25-200-0.37-30, Inper) to a rotary joint (FRJ_1 × 2_FC-2FC, Doric Lenses), then with an DPSS laser (MBL-F-473–200 mW, Shanghai Dream Lasers). Laser power was 15–20 mW measured at the distal tip of patch cord. Master-8 Pulse Stimulator (Advanced Medical Physics Instrumentation) was used to control the parameter of laser stimulation. The 30-min stimulation session consisted of 60 alternating 15-s oDAS-on and 15-s oDAS-off epochs. Each oDAS-on epoch consisted of 30 bursts (duration: 250 ms, interval: 250 ms). Each burst comprises 5-pulse of laser (10 ms pulse at a frequency of 20 Hz).

### Cocaine injection

Mice were injected intraperitoneally (i.p.) with 15 mg kg^−1^ cocaine (dissolved in saline; injection volume: 10 mL kg^−1^). For the cocaine challenge, animals were injected immediately after the free period, placed back in test apparatus, and recordings were started.

### Locomotion sensitization

Locomotor activity was measured as the distance traveled in a grey opaque circular corridor (inner diameter: 10 cm, height: 24 cm). The test apparatus with a transparent bottom was placed above a camera (FLIR, framerate: 40 Hz) and recorded by video tracking system (CinePlex, Plexon). Tests were performed during the light phase of the light/dark cycle.

Animals were placed in the circular corridor for 3 daily sessions of 30 minutes for habituation. For the next 5 days and for the challenge day 15, they were first placed in the arena for 30 minutes (free period) and the stimulation started for another period of 30 minutes (stimulation period).

### Video analysis

From the videos at the framerate of 40 Hz, the initial pose estimate was acquired through Plexon, and then refined using a custom MATLAB script. The center of the arena was computed for each video from the leftmost, rightmost, topmost, and bottommost coordinate on the trajectory of the animal. This estimate was also used to convert the distances from pixel space to centimeters. The trajectory was interpolated over absent values and/or coordinates detected outside the arena using shape-preserving piecewise cubic interpolation. Finally, the trajectory was smoothed with a moving average filter of width 5. Data were further analyzed with Microsoft excel 16.54, Igor Pro 6.34 and GraphPad prism 9.

## Bibliography

1. Robinson, T.E., and Berridge, K.C. (2008). Review. The incentive sensitization theory of addiction: some current issues. Philosophical transactions of the Royal Society of London. Series B, Biological sciences 363, 3137–3146. 10.1098/rstb.2008.0093.

2. Herve, D., Levi-Strauss, M., Marey-Semper, I., Verney, C., Tassin, J., Glowinski, J., and Girault, J. (1993). G(olf) and Gs in rat basal ganglia: possible involvement of G(olf) in the coupling of dopamine D1 receptor with adenylyl cyclase. J. Neurosci. 13, 2237–2248. 10.1523/JNEUROSCI.13-05-02237.1993.

3. Girault, J.-A. (2012). Signaling in striatal neurons: the phosphoproteins of reward, addiction, and dyskinesia. Progress in molecular biology and translational science 106, 33– 62. 10.1016/B978-0-12-396456-4.00006-7.

4. Bertran-Gonzalez, J., Bosch, C., Maroteaux, M., Matamales, M., Hervé, D., Valjent, E., and Girault, J.-A. (2008). Opposing patterns of signaling activation in dopamine D1 and D2 receptor-expressing striatal neurons in response to cocaine and haloperidol. The Journal of neuroscience : the official journal of the Society for Neuroscience 28, 5671–5685. 10.1523/JNEUROSCI.1039-08.2008.

5. Pascoli, V., Besnard, A., Hervé, D., Pagès, C., Heck, N., Girault, J.-A., Caboche, J., and Vanhoutte, P. (2011). Cyclic adenosine monophosphate–Independent tyrosine phosphorylation of NR2B mediates cocaine-induced extracellular signal-regulated kinase activation. Biological Psychiatry 69, 218–227. 10.1016/j.biopsych.2010.08.031.

6. Valjent, E., Pagès, C., Hervé, D., Girault, J., and Caboche, J. (2004). Addictive and non-addictive drugs induce distinct and specific patterns of ERK activation in mouse brain. Eur J of Neuroscience 19, 1826–1836. 10.1111/j.1460-9568.2004.03278.x.

7. Valjent, E., Corvol, J.-C., Trzaskos, J., Girault, J.-A., and Hervé, D. (2006). Role of the ERK pathway in psychostimulant-induced locomotor sensitization. BMC Neurosci 7, 20. 10.1186/1471-2202-7-20.

8. Pascoli, V., Turiault, M., and Lüscher, C. (2012). Reversal of cocaine-evoked synaptic potentiation resets drug-induced adaptive behaviour. 481, 71–75. 10.1038/nature10709.

9. Creed, M.C. (2017). Toward a targeted treatment for addiction. Science (New York, N.Y.) 357, 464–465. 10.1126/science.aao1197.

10. Pascoli, V., Terrier, J., Espallergues, J., Valjent, E., O’Connor, E.C., and Lüscher, C. (2014). Contrasting forms of cocaine-evoked plasticity control components of relapse. 509, 459– 464. 10.1038/nature13257.

11. Ungless, M.A., Whistler, J.L., Malenka, R.C., and Bonci, A. (2001). Single cocaine exposure in vivo induces long-term potentiation in dopamine neurons. 411, 583–587. 10.1038/35079077.

12. Bellone, C., and Lüscher, C. (2006). Cocaine triggered AMPA receptor redistribution is reversed in vivo by mGluR-dependent long-term depression. Nature neuroscience 9, 636– 641. 10.1038/nn1682.

13. Pascoli, V., Terrier, J., Hiver, A., and Lüscher, C. (2015). Sufficiency of mesolimbic dopamine neuron stimulation for the progression to addiction. Neuron 88, 1054–1066.

14. Chaudun, F., Python, L., Liu, Y., Hiver, A., Cand, J., Kieffer, B.L., Valjent, E., and Lüscher, C. (2024). Distinct µ-opioid ensembles trigger positive and negative fentanyl reinforcement. Nature 630, 141–148. 10.1038/s41586-024-07440-x.

15. Robinson, T.E., and Berridge, K.C. (2026). Can the incentive-sensitization theory of addiction incorporate addiction to opioid drugs? Psychopharmacology (Berl). 10.1007/s00213-025-07001-8.

